# The zinc finger protein CLAMP promotes long-range chromatin interactions that mediate dosage compensation of the *Drosophila* male X-chromosome

**DOI:** 10.1101/2020.11.02.365122

**Authors:** William Jordan, Erica Larschan

## Abstract

*Drosophila* dosage compensation is an important model system for defining how active chromatin domains are formed. The Male-specific lethal dosage compensation complex (MSLc) increases transcript levels of genes along the length of the single male X-chromosome to equalize with that on the two female X-chromosomes. The strongest binding sites for MSLc cluster together in three-dimensional space independent of MSLc because clustering occurs in both sexes. CLAMP, a non-sex specific, ubiquitous zinc finger protein, binds synergistically with MSLc to enrich the occupancy of both factors on the male X-chromosome. Here, we demonstrate that CLAMP promotes the observed clustering of MSLc bindings sites. Genome-wide, CLAMP promotes interactions between active chromatin regions. Moreover, the X-enriched CLAMP protein more strongly promotes longer-range interactions on the X-chromosome than autosomes. Genome-wide, CLAMP promotes interactions between active chromatin regions together with other insulator proteins. Overall, we define how long-range interactions which are modulated by a locally enriched ubiquitous transcription factor promote hyper-activation of the X-chromosome to mediate dosage compensation.

## Introduction

Three-dimensional chromatin domains are important for coordinating gene regulation. Recent work has provided new insight into how silent chromatin domains are formed, for example, through phase separation^1,2^, but less is understood regarding the formation of hyper-active chromatin domains^3^. Dosage compensation in *Drosophila* provides one of the few model systems for studying the formation of a large hyper-active chromatin domain: approximately one thousand active genes along the length of the single male X-chromosome are coordinately upregulated 2 fold^4–6^.

In heterogametic species, dosage compensation is essential to correct transcriptional imbalance of X-linked genes between the sexes^7^ and to correct for dosage imbalance between the single X-chromosome and paired autosomes. Diverse dosage compensation mechanisms have evolved across species, but an essential conserved step is distinguishing the X-chromosome from autosomes for specific regulation.

In *Drosophila,* the male-specific lethal complex (MSLc) forms only in males and is responsible for increasing transcript levels of X-linked genes along the length of the single male X-chromosome 1.4 fold, helping to equalize gene expression with that of females^4–6^. MSLc consists of five proteins MSL1, MSL2, MSL3, maleless (MLE)^8–10^, males absent on the first (MOF)^11^, and one of two functionally redundant long non-coding RNAs known as RNA on the X 1 and 2 *(roX1* and *roX2)*^12,13^. MSLc first targets X-linked genomic elements known as “high-affinity” (HAS) or “chromatin entry” sites (CES), which also include the *roX* loci^12,14–16^.

Within CES, MSLc is recruited to GA-rich 21-bp elements known as MSL recognition elements (MREs)^14^ Accumulation of MRE sequences on the X-chromosome occurred by expansion of GA-rich sequences and transposon insertion^17,18^. However, MRE sequences are not X-chromosome specific and are only approximately two-fold enriched on the X-chromosome compared with autosomes^14^, suggesting they are not sufficient for X-chromosome targeting. Although the MSL2 component of MSLc has a low affinity for MREs, MSL complex requires synergy with an essential, non sex-specific, zinc finger adapter protein known as chromatin-linked adapter for MSL proteins (CLAMP) in order to stabilize its binding to MREs^19–21^.

Synergy between CLAMP and MSLc, which has been demonstrated both *in vivo* and *in vitro*^20,21^, enhances the occupancy of both factors on the male X-chromosome. Maternally deposited CLAMP is present on chromatin throughout the genome before MSLc assembles at the maternal-zygotic transition^22–24^ and regulates chromatin accessibility of the X-chromosome and transcription of X-linked genes^19,25^. Therefore, it is likely that CLAMP functions as an early transcription factor to enhance X-chromosome accessibility and promote MSLc targeting.

After initial targeting to CES by CLAMP, MSLc generates a hyper-active chromatin domain by localizing to the bodies of active genes and increasing their transcript levels through modulating transcription elongation^14,15,25–27^. MSLc was hypothesized to take advantage of pre-existing three-dimensional chromatin organization to target the X-chromosome^28,29^. Chromosome conformation capture techniques have demonstrated that CES cluster three-dimensionally in both males and females and form long-range interactions with other X-linked active chromatin regions within the nucleus independent of MSLc^28,29^. However, the mechanism by which CES cluster remained unknown.

We hypothesized that CLAMP promotes clustering of CES based on the following lines of evidence: 1) In contrast to MSLc, CLAMP is required to globally increase the accessibility of the entire male X-chromosome^25^; 2) CLAMP is part of two insulator protein complexes (Kaye et al., 2017; Bag et al., 2019), acts as an insulator protein in several functional assays^30^, and promotes recruitment of the CP190 insulator protein^30^. Insulator proteins mediate chromatin interactions across the genome to regulate specialized chromatin domains throughout development^31–35^. However, it was not known whether CLAMP regulates the formation of three-dimensional interactions within the genome.

We used genome-wide chromosome conformation capture (Hi-C) analysis complemented by circular chromosome conformation capture with high-throughput sequencing (4C-seq) to test the hypothesis that CLAMP regulates clustering of CES and three-dimensional organization of the X-chromosome. We discovered that CLAMP promotes long-range interactions on the male X-chromosome more strongly than on autosomes. Furthermore, we demonstrate that CLAMP primarily promotes long-range interactions within active chromatin regions, including CES. We also show that enrichment of several insulator proteins is increased at loci where CLAMP regulates genomic interactions. Overall, we demonstrate that the X-enriched CLAMP protein regulates long-range three-dimensional interactions between CES to target MSLc to the male X-chromosome. Synergy between CLAMP and MSLc^20,21^ increases the occupancy of both factors to specifically hyper-activate approximately one thousand X-linked genes in males.

## Results

### CLAMP promotes long-range three-dimension interactions on the X-chromosome more strongly than on autosomes

In order to understand how CLAMP regulates the three-dimensional organization of the genome, we performed *in situ* chromosome conformation capture with high-throughput sequencing (*in situ* Hi-C)^36^ using HindIII, a 6-bp cutter restriction enzyme, in *Drosophila* male Schneider’s line 2 (S2)^37^ cultured cells after validated RNAi depletion of either *gfp* (control) or *clamp*^4,19,20^. We performed two biological replicates for each experimental condition. We confirmed depletion of CLAMP protein after *clamp* RNAi by Western blot (Fig. S1A) and used GenomeDISCO^38^ to determine replicate concordance. Reproducibility between replicates was high (Figure S1B).

We visualized our Hi-C interaction maps by combining replicates for each condition (Fig. S1C; Table S1). To compare the *clamp* RNAi and control,*gfp* RNAi contact maps, we also generated a differential interaction map (Fig. S1C). In the differential map for the two conditions (*clamp/gfp* RNAi), we observed an increase in interaction frequency (red) directly along the diagonal (i.e. shorter-range interactions). Many of the more pronounced off-diagonal differences are in regions proximal to centromeres, which will be discussed later. In general, we observe decreased interaction frequency (blue) moving away from the diagonal (i.e. longer-range interactions) across all chromosomes when CLAMP is depleted and this decrease is more widespread on the X-chromosome compared with autosomes.

To quantify differences between our *gfp* and *clamp* RNAi interaction matrices, we calculated the log_2_ ratio of distal to local interactions (DLR)^39^ (Fig. 1, Fig. S2A). We defined local interactions as those that span less than 250 kb and distal as interactions that span more than 250 kb based on the average size of a TAD in *Drosophila.* After depletion of CLAMP, there is a change in the ratio between long-range and short-range interactions that is different on the X-chromosome compared with autosomes. The X-enriched decrease in the DLR after *clamp* RNAi is also observed when DLR is measured using a local vs distal cutoff of 100 kb instead of 250 kb (Fig. S2B). Moreover, this X-enriched change in three-dimensional interactions (Fig. 1, Fig. S2A,B) is consistent with previous MNase accessibility analysis demonstrating that global chromatin accessibility of the male X-chromosome but not autosomes decreases after *clamp* RNAi^25^. Therefore, our Hi-C data support a model in which CLAMP alters the three-dimensional organization of the male X-chromosome more than autosomes.

**Figure 1.**
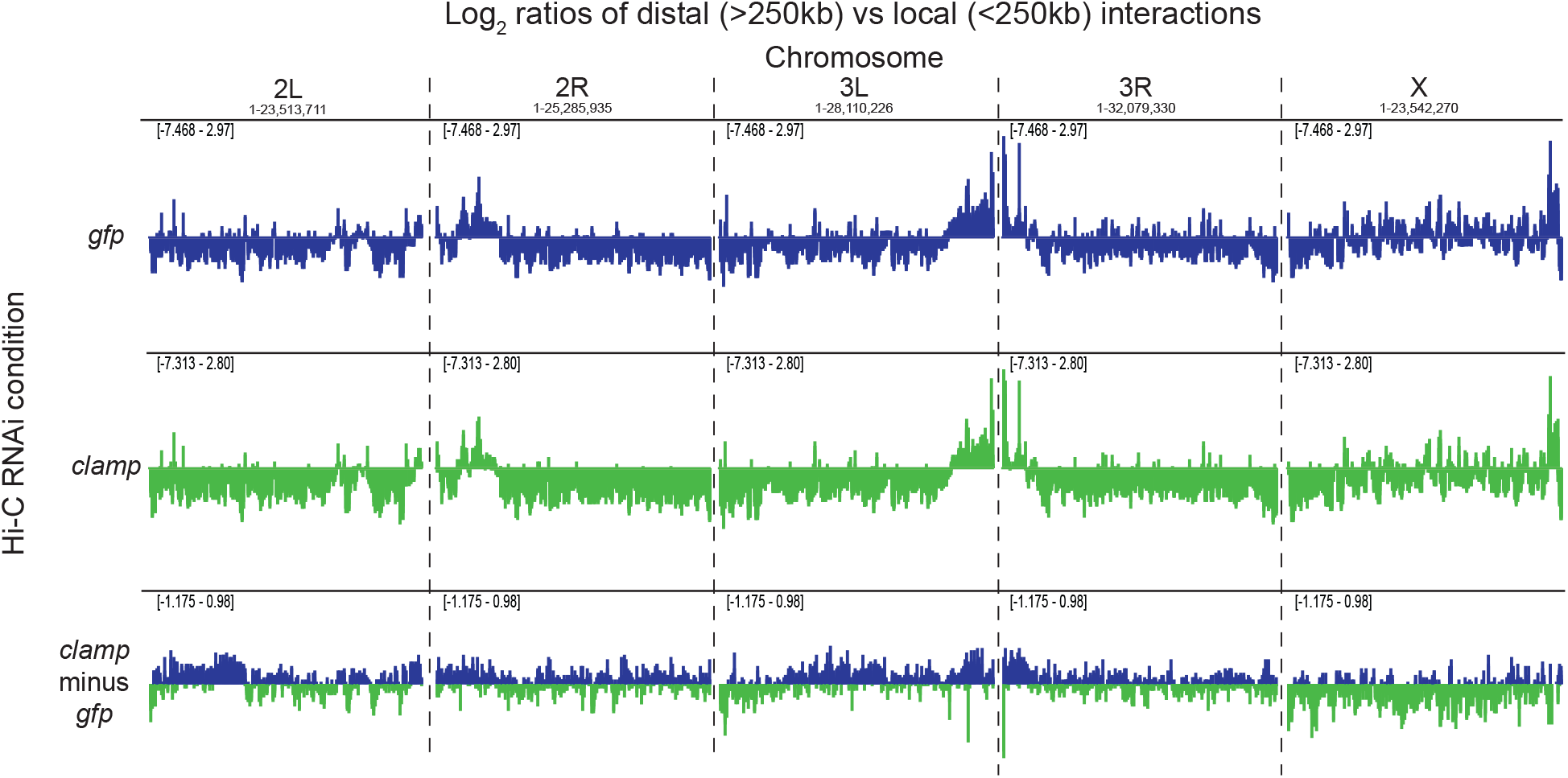
CLAMP regulates the length span of genomic interactions on the male X-chromosome. Per chromosome distal vs local ratio (DLR) for *gfp* RNAi (blue), *clamp* RNAi (green), and *clamp* vs *gfp* RNAi (bottom). For the *clamp* vs *gfp* RNAi comparison, a positive number (blue) indicates the ratio of distal vs. local interactions becomes higher following *clamp* RNAi. A negative number (green) indicates the ratio of distal vs. local interactions becomes lower following *clamp* RNAi.

Previous Hi-C studies in multiple cell lines and embryos found that the strongest MSLc binding sites (CES) interact with each other and other genomic regions along the X-chromosome more frequently than expected by chance^28,29^. To confirm these observations within our own Hi-C data, we investigated whether CES interact frequently with other genomic locations. We used Fit-Hi-C^40^ to determine high-confidence intra-chromosomal contacts (see methods) from our control *gfp* RNAi Hi-C maps (Fig. S1D; Table S3). Consistent with previous reports^28,29^, approximately 60% (3,090) of the high-confidence X-chromosome interactions identified at 20 kb resolution involve CES; 45% would be expected by chance, even when restricting our analysis to only active chromatin regions as controls (p = 5.7e-107, chi-square test) (Fig. S1E). Therefore, our data are consistent with prior reports that CES interact more frequently with other regions of the genome than expected by chance. Moreover, we demonstrate that CES interact more frequently with other regions of the genome even when compared with other active chromatin regions that are known to cluster together.

To more quantitatively define specific regions throughout the genome where CLAMP regulates three-dimensional interactions, we compared intra-chromosomal interaction frequencies after *clamp* RNAi with those after *gfp* RNAi using diffHic^41^. DiffHic uses edgeR to model biological variability between replicates and perform differential analysis between conditions^41,42^. We found 2,552 significantly (FDR < 0.05) differential interactions (DIs) at 30 kb resolution (Fig. S3A; Table S4). We classified these DIs by the directionality of their log_2_ ratio. Notches on all box plots represent 95% confidence intervals around the median line; whiskers represent 1.5 IQR (inter-quartile range) and outliers have been omitted. Interactions that decrease in contact probability after *clamp* RNAi are defined as CLAMP-promoted (55%), whereas interactions that increase in contact probability after *clamp* RNAi are defined as CLAMP-repressed (45%). The log2 ratio was significantly larger for CLAMP-promoted X-linked interactions (Fig. 2A; Fig. S3B) than other remaining interaction types throughout the genome. Therefore, CLAMP more strongly promotes three-dimensional interactions on the X-chromosome compared to on autosomes.To define the properties of DIs mediated by CLAMP, we measured the genomic distance between DI anchors. The linear distance spanned by CLAMP-promoted X-linked interactions is significantly longer than those that CLAMP promotes on autosomes (Fig. 2B; Fig. S3C). In contrast, the length span of interactions repressed by CLAMP is significantly shorter on the X-chromosome than on autosomes (Fig. 2B; Fig. S3C), consistent with the X-biased decrease in DLR after depleting CLAMP (Fig. 1, Fig S2A, B). Therefore, CLAMP promotes long-range interactions on the X-chromosome. We also measured the proximity of CLAMP-promoted and CLAMP-repressed DI anchors to CES and two randomized classes of sites within regions of either active or inactive chromatin based on the 9-state chromatin state model for S2 cells^43^ (Fig. 2C). We found that CLAMP-promoted regions are closer to active chromatin regions than CLAMP-repressed regions. In contrast, CLAMP-repressed regions are closer to inactive chromatin than active chromatin regions. Also, CES are in closer proximity to CLAMP promoted DI anchors and more distal from CLAMP-repressed DI anchors than other active chromatin regions (Fig. 2C). Overall, CLAMP promotes long-range contacts on the X-chromosome that enhance three-dimensional interactions involving active chromatin and CES.

**Figure 2.**
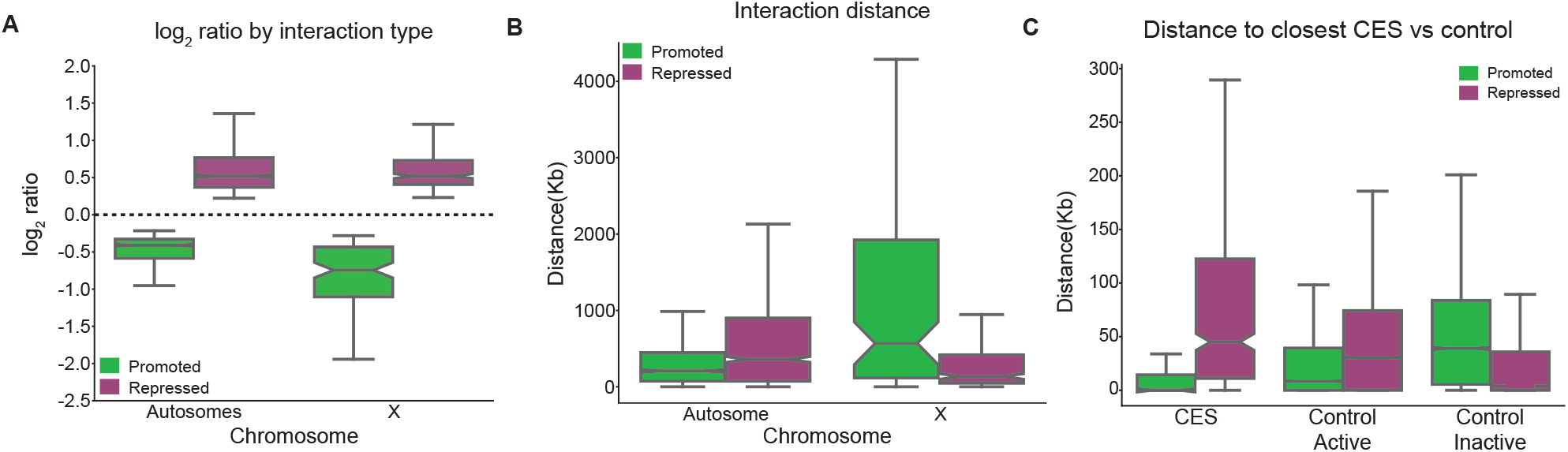
CLAMP promotes the formation of longer-range contacts more strongly on the X-chromosome than autosomes. **A.** log_2_ ratios per chromosome of interactions that are weakened after *clamp* RNAi (CLAMP-promoted) or strengthen (CLAMP-repressed) (Source data provided in Table S4). **B.** Distribution of distances between DI anchors. On the autosomes, CLAMP-promoted interactions are shorter-range compared to autosomal CLAMP-repressed interactions. On the X-chromosome however CLAMP-promoted interactions are much longer-range than CLAMP-repressed (Source data provided in Table S4). **C.** Distribution of distances to nearest CES or control region for CLAMP-promoted and CLAMP-repressed interactions. For all box and whisker plots, the 95% confidence interval is shown with a notch around the median line; whiskers represent 1.5 IQR, outliers have been omitted. (**a – b** CLAMP promoted interactions: autosomes n = 1055, X n = 103; CLAMP repressed interactions: autosomes n = 1142, X n = 252. **c** Promoted n = 206, repressed n = 504; distribution of controls obtained by 100 permutations of randomly shuffling CES (see methods; Source data provided as a Source Data file).

### CLAMP promotes three-dimensional interactions at active chromatin regions including CES

Next, we defined the relationship between DIs that are regulated by CLAMP and the enrichment of the CLAMP protein. First, we used available CLAMP ChIP-seq data^44^ from S2 cells to generate a high-confidence list of CLAMP peaks and average profiles of CLAMP peak enrichment at DI anchors (see methods). We found that CLAMP-promoted DI anchors more frequently contain CLAMP peaks than CLAMP-repressed DI anchors on both the X-chromosome and autosomes (Fig. 4A). Therefore, CLAMP-promoted contacts are more likely to be directly linked to CLAMP function than CLAMP-repressed contacts. Furthermore, we found that 60% of CLAMP-promoted DI anchors on the X-chromosome are occupied by CLAMP, which is greater than CLAMP occupancy at CLAMP-repressed anchors on the X-chromosome or CLAMP-promoted and CLAMP-repressed anchors on autosomes. While CLAMP frequently binds to DI anchors, the ChIP-seq occupancy of CLAMP is not more enriched at these genomic locations where CLAMP regulates three-dimensional genomic interactions compared to CLAMP sites that do not occur at DI anchors. Therefore, we hypothesize that the presence of additional cofactors within active and/or inactive chromatin modulate the ability of CLAMP to influence three-dimensional interactions.

To test this hypothesis, we first measured the chromatin states, as defined by the *Drosophila* 9-state chromatin model^43^, that are present at CLAMP-promoted and CLAMP-repressed DI anchors. We found that across all chromosomes, 60.5% of the chromatin states present within CLAMP-promoted DI anchors represent active chromatin and the remaining 39.5% of chromatin states represent inactive chromatin (Fig S3D). In contrast, 70% of the chromatin states present within CLAMP-repressed DI anchors represent inactive states, while the remaining 30% of chromatin states represent active chromatin across all chromosomes (Fig S3E). Additionally, at CLAMP-repressed DI anchors, there is an enrichment of the chromatin state corresponding to pericentromeric heterochromatin (Fig. S3E) which correlates with our visual observation that many pronounced off-diagonal changes in our differential Hi-C maps occur in regions proximal to centromeres (Fig. S1C).

To further quantify the relationship between DI anchors and chromatin states, we computed Jaccard similarity coefficients as ratios ranging from 0 to 1^45^: the larger the Jaccard coefficient (i.e. closer to 1), the more similar two sets of genomic regions are to each other. CLAMP-promoted DI anchors have more similarity to active chromatin states (Jaccard coefficient: 0.328) than inactive chromatin states (Jaccard coefficient: 0.220). In contrast, CLAMP-repressed DI anchors have more similarity to inactive chromatin states (Jaccard coefficient: 0.297) than active chromatin states (Jaccard coefficient: 0.090). Therefore, CLAMP primarily promotes interactions within active regions; interactions that form after *clamp* RNAi are often within inactive regions of the genome. Moreover, the ability of CLAMP to promote genomic interactions within active chromatin regions (Fig. S3D; Fig. S3E) is consistent with its ability to activate gene expression and open chromatin on the active male X-chromosome more frequently than on autosomes^25^.

### CLAMP and MSLc both mediate three-dimensional interactions at CES

We and others previously reported that CLAMP and MSLc function synergistically to target each other to the X-chromosome and increase occupancy of both factors *in vivo* and *in vitro*^20,21^. In addition, MSLc modulates chromatin accessibility specifically within CES, in contrast to CLAMP which not only functions at CES but also enhances chromatin accessibility of the entire male X-chromosome^25,28^. Furthermore, long-range interactions at several CES occur in an MSL2-dependent manner^29,46^. Therefore, we hypothesized that both CLAMP and MSL complex function to regulate three-dimensional interactions at CES. Therefore, we also processed and performed differential interaction analysis with available Hi-C data (GSE58821) from replicated dilution Hi-C experiments that compare RNAi depletion of two MSLc components (MSL2 and MSL3) with a matched control *(gfp* RNAi) experiment^28^ (Fig S1B). In contrast to our *clamp* RNAi experiments, we did not identify significant DIs following *msl2* RNAi, and *msl3* RNAi resulted in only 1 significant DI when compared to matched *gfp* controls (Table S4). Therefore, MSLc does not modulate three-dimensional interactions detectable by dilution Hi-C consistent with previous reports (Ramirez et al., 2015).

To validate our Hi-C findings that CLAMP regulates three-dimensional organization and generate a higher resolution subset of DIs, we performed circularized chromosome conformation capture with high-throughput sequencing (4C-seq) at four CES in close proximity to regions containing many DIs identified by Hi-C. In addition, we assessed the function of MSL2 and the GAF protein (encoded by the *trl* gene), which is a GA-binding zinc-finger protein similar to CLAMP. CLAMP and GAF are present in the same insulator complex^44,47–49^. However, in contrast to CLAMP, GAF has only a modest function in MSLc recruitment and is depleted within CES because CLAMP outcompetes GAF for binding to the long GA-rich sequences that are present within CES^44,50^. Therefore, we hypothesized that CLAMP and MSL2 but not GAF regulate three-dimensional interactions at CES.

To test this hypothesis, we performed 4C-seq experiments in biological duplicate in *Drosophila* S2 cells following previously validated RNAi depletion of *gfp* (control), *msl2, clamp* and *trl*^20,44^ and confirmed depletions by qRT-PCR (Fig. S4A; Table S1; Table S2). We identified high frequency *cis*-interacting regions and performed differential interaction analysis for each viewpoint using the 4C-ker pipeline^51^ (Fig. S4). Consistent with our Hi-C analysis, we found that 59% of high-confidence *cis*-interactions link our four CES viewpoints to regions that also contain a CES (Fig S4B; Table S3). After visualizing differential interactions for each viewpoint (Table S4), we pooled all identified DI anchors from each 4C-seq viewpoint to increase the number of DIs for further analysis.

Next, we compared the number of DIs obtained after *clamp* and *tr1*RNAi. We identified 199 total DIs after *clamp* RNAi and only 17 DIs after *trl* RNAi (Fig S4C). Therefore, even though both CLAMP and GAF are part of the same insulator complex^49^, CLAMP has a stronger role than GAF in modulating three-dimensional interactions involving CES. These data are consistent with an enrichment of CLAMP versus GAF at CES^44^ Next, we measured chromatin state occurrence within CLAMP-promoted and CLAMP-repressed 4C-seq DI anchors and found similar chromatin states to those identified in our differential Hi-C analysis (Fig. S4F and S3E-F). Therefore, our 4C-seq data demonstrate that CLAMP has a stronger role than GAF in regulating three-dimensional interactions at CES and validate a role for CLAMP in promoting contacts between active chromatin regions.

In addition, we measured the role of CLAMP and MSL2 in regulating three-dimensional interactions at CES by comparing our 4C-seq data after *clamp* and *msl2* RNAi treatments. In contrast to our differential Hi-C interaction analysis which did not identify any DIs after *msl2* RNAi, we observed 285 DIs after *msl2* RNAi from our CES viewpoints, consistent with prior findings^29,46^ (Fig S4C). The discrepancy between the Hi-C and 4C-seq analyses for MSL2 may be due to the dilution Hi-C technique performed by Ramirez et al. which contains more technical noise because there is higher potential for spurious contacts during in-solution ligation compared with *in situ* ligation^36,52,53^. Furthermore, the Hi-C and 4C techniques have very different resolutions. We also measured X-specific chromatin state occurrence for 4C-seq DI anchors identified after *msl2* RNAi and we find similar chromatin states to those identified at CLAMP-dependent 4C-seq DI anchors (Fig. S4E-F). Overall, both CLAMP and MSL2 regulate threedimensional interactions at CES, consistent with prior observations that CLAMP and MSLc modulate chromatin accessibility^25,28^ and promote each other’s occupancy^20,21^ locally at CES.

To further define the relationship between CLAMP and MSLc in mediating three-dimensional interactions, we integrated our Hi-C and 4C data with previously-generated chromatin accessibility data from a Micrococcal Nuclease (MNase)-seq experiment performed in S2 cells under the same RNAi conditions (GSE99894)^25^. In MNase-seq experiments, a MNase accessibility score (MACC) greater than zero indicates that a region of chromatin is relatively accessible while a MACC score less than zero indicates that a region is relatively inaccessible compared to the average genomic accessibility. We found that CLAMP-promoted and CLAMP-repressed DIs from both Hi-C and 4C occur within regions where the chromatin is relatively accessible (MACC >0) under control *gfp* and *msl2* RNAi conditions (Fig. 3A, B). However, significant decreases in chromatin accessibility after *clamp* RNAi, are observed at CLAMP-promoted and CLAMP-repressed DI anchors on the X-chromosome and CLAMP-repressed DIs on autosomes (Fig. 3A, B). Therefore, CLAMP regulates chromatin accessibility at regions where it regulates three-dimensional interactions.

**Figure 3.**
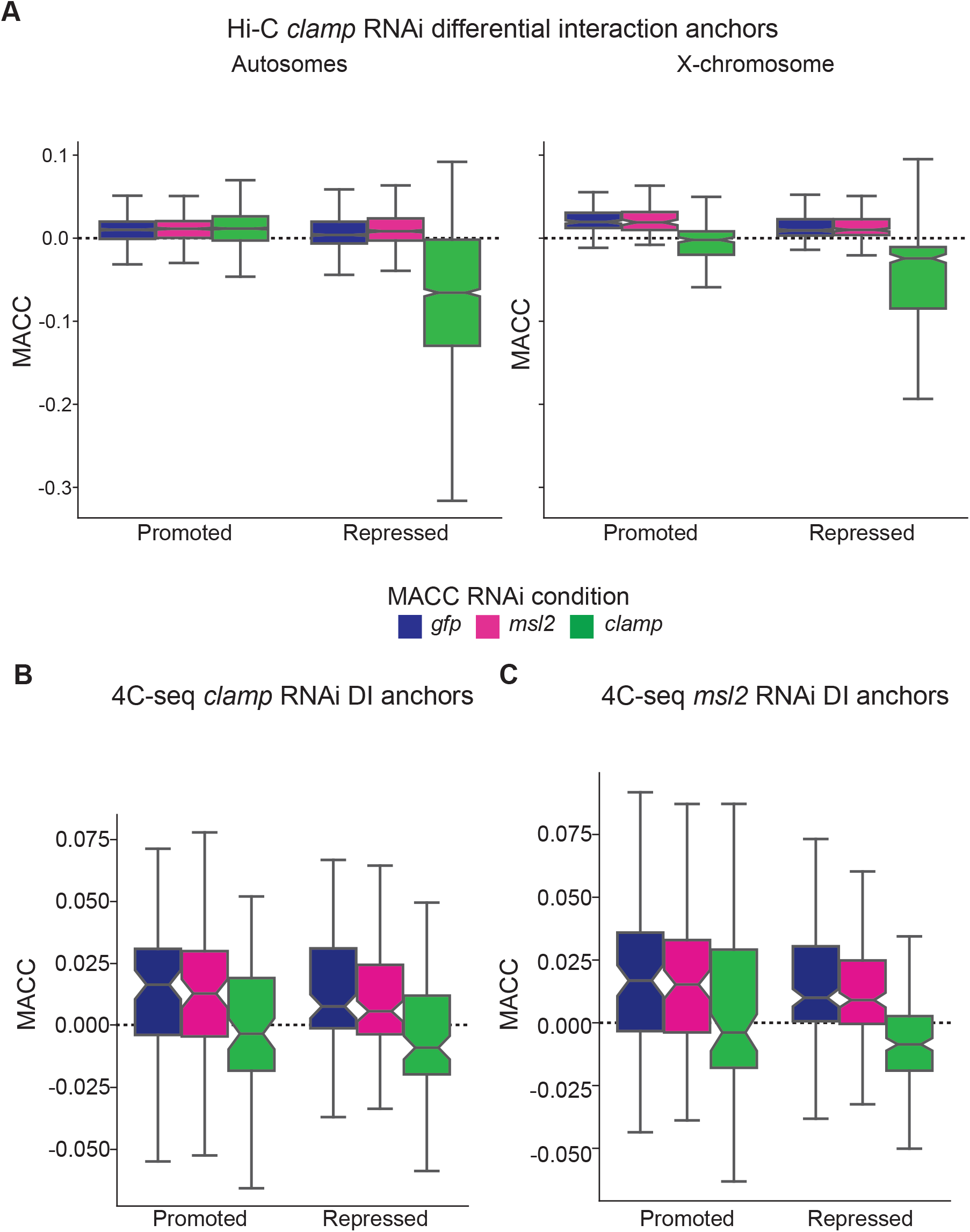
CLAMP regulation of the three-dimensional organization is linked to its role in altering chromatin accessibility. **A.** Distribution of chromatin accessibility MACC values within CLAMP Hi-C DI anchors as measured when MACC was previously calculated under RNAi conditions of *gfp* (blue), *msl2* (pink) and *clamp* (green). A positive MACC value indicates accessible chromatin whereas a negative MACC value indicates inaccessible chromatin^25^. On the autosomes, CLAMP promoted interactions occur in regions that are accessible under all three RNAi conditions. In contrast, autosomal CLAMP-repressed interactions occur in regions that have lowered chromatin accessibility after *clamp* RNAi (Promoted interactions: *gfp* RNAi n = 2104, *msl2* RNAi n = 1052, *clamp* RNAi n = 2104; Repressed interactions: *gfp* RNAi n = 1968, *msl2* RNAi n = 984, *clamp* RNAi n = 1968). On the X-chromosome both CLAMP-promoted and repressed interactions occur in regions which are accessible after *gfp* and *msl2* RNAi but become inaccessible after *clamp* RNAi (Promoted interactions: *gfp* RNAi n = 196, *msl2* RNAi n = 98, *clamp* RNAi n = 196; Repressed interactions: *gfp* RNAi n = 466, *<u>msl2</u>* RNAi n = 233, *clamp* RNAi n = 466). **B.** Distribution of MACC values within CLAMP 4C-seq DI anchors. CLAMP 4C-seq identified DIs occur in regions which have significantly lowered chromatin accessibility after *clamp* RNAi (green) compared to control *gfp* RNAi (blue) (Promoted interactions: *gfp* RNAi n = 87, *msl2* RNAi n = 87, *clamp* RNAi n = 87; Repressed interactions: *gfp* RNAi n = 110, *msl2* RNAi n = 110, *clamp* RNAi n = 110).. No significant changes are observed for these CLAMP 4C-seq identified DI regions are observed following *msl2* RNAi (pink). **C.** Distribution of MACC values within MSL2 4C-seq DI anchors. MSL2 4C-seq identified DIs occur in regions that do not have significant changes in chromatin accessibility after *msl2* RNAi (pink) compared to control *gfp* RNAi (blue). However, chromatin accessibility is significantly decreased following *clamp* RNAi (green) (Promoted interactions: *gfp* RNAi n = 101, *msl2* RNAi n = 101, *clamp* RNAi n = 101; Repressed interactions: *gfp* RNAi n = 181, *msl2* RNAi n = 181, *clamp* RNAi n = 181). For all box and whisker plots, the 95% confidence interval is shown with a notch around the median line; whiskers represent 1.5 IQR, outliers have been omitted. (Source data provided as a Source Data file).

Next, we asked whether CLAMP or MSL2 regulates chromatin accessibility at sites where MSL2 regulates three-dimensional interactions. We determined that MSL2-dependent 4C-seq DIs show significant decreases in chromatin accessibility after *clamp* RNAi and modest but not statistically significant changes after *msl2* RNAi (Fig. 3C). Therefore, CLAMP but not MSL2 regulates chromatin accessibility at regions where MSL2 regulates three-dimensional interactions, consistent with the synergy between the two factors. Overall, the function of CLAMP in modulating three-dimensional interactions is linked to its role in altering chromatin accessibility.

### Insulator proteins have differential occupancy at genomic locations at which CLAMP regulates three-dimensional interactions

CLAMP has been physically and functionally linked with two different insulator complexes containing either the insulator proteins GAF^49^ or Su(Hw)^30^. Therefore, we hypothesized that the function of CLAMP in regulating three-dimensional interactions is mediated by differential occupancy of insulator proteins at DI anchors. To test this hypothesis, we generated high confidence peaks and average profiles for insulator proteins from the following publicly available ChIP-seq data generated in S2 cells for the insulator proteins GAF (GSE107059), CP190, Su(Hw), Mod(mdg4) and dCTCF (GSE41354). We restricted our analysis to CLAMP-dependent DIs identified by Hi-C in order to make genome-wide comparisons and compared ChIP-seq enrichment of each factor at peaks that occur within DI anchors (DIs) to enrichment at the remaining set of peaks that occur outside of DI anchors (non-DIs).

Overall, we observed enhanced occupancy of all insulator proteins at genomic anchor points where CLAMP regulates three dimensional interactions (DIs) compared with regions where it does not regulate three-dimensional interactions (non-DIs). Furthermore, we discovered the following differences in factor enrichment: 1) Overall the enrichment patterns of GAF are similar to that of CLAMP, which correlates with their presence in the same insulator complex^44^ Also, GAF enrichment as measured by ChIP-seq is depleted at CLAMP-promoted DI anchors compared with those outside of DI anchors or CLAMP-repressed anchors (Fig. 4B). 2) Su(Hw) is bound to fewer CLAMP-repressed DI anchors on autosomes compared with those on the X-chromosome, although its average enrichment at bound sites is similar. (Fig. 4C). 3) Mod(mdg4), dCTCF and CP190 are all more frequently bound at CLAMP-promoted DI anchors. However, their enrichment is higher at CLAMP-repressed DI anchor sites versus CLAMP-promoted anchors. Furthermore, CP190 binds to a larger majority of CLAMP-promoted DIs than any of the other factors tested consistent with a previously reported role for CLAMP in modulating CP190 recruitment^29^ (Fig. 4D, E, F).

**Figure 4.**
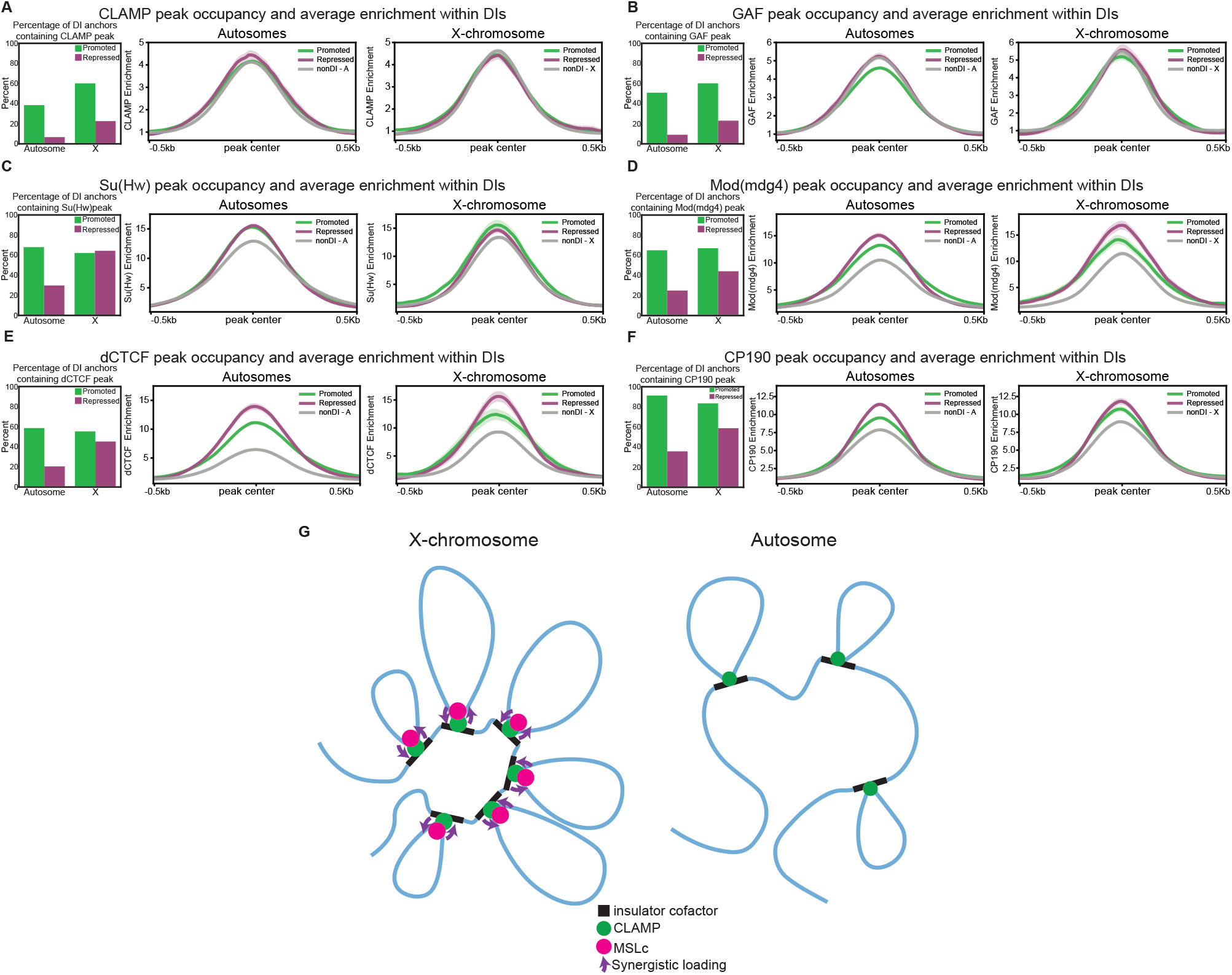
Differential occupancy of insulator proteins occurs at loci where CLAMP promotes or represses the formation of three-dimensional interactions on the X-chromosome and autosomes. **A.** Left: Percentage of DI anchors containing a high confidence CLAMP ChIP-seq peak. Right: average fold enrichment over input for CLAMP at CLAMP peaks within CLAMP-promoted DI anchors (green) and CLAMP-repressed DI anchors (purple) versus the remaining set of peaks falling outside of DIs (gray). Light shading on average profiles represents standard error. The same analysis was performed for GAF (**B**), Su(Hw) (**C**), Mod(mdg4) (**D**), dCTCF (**E**), and CP190 **(F**) (Source data provided as a Source Data file). CLAMP promotes long-range interactions on the X-chromosome that are responsible for the clustering of CES. Synergy between CLAMP and MSL complex increases the local concentration of both factors on the male X-chromosome. On autosomes, the local concentration of CLAMP is not as high due to the lack of synergy with MSL complex **(G).** Model for function of CLAMP in three-dimensional organization. ^20^ which reduces the number of long-range interactions that are mediated by CLAMP.

Overall, we observe different patterns of insulator protein occupancy within CLAMP-promoted and CLAMP-repressed DI anchors, compared to peak locations where we do not detect CLAMP-dependent three-dimensional interactions (non-DIs). CLAMP is also a member of two different insulator protein complexes^29,43^. Therefore, CLAMP may modulate recruitment and/or function of insulator protein complexes to regulate three-dimensional interactions at DI anchors. In fact, very recent work demonstrated that CLAMP regulates the occupancy of the CP190 protein^30^.

## Discussion

Overall, our data provide key insight into how the *Drosophila* male X-chromosome forms a specific hyper-active chromatin domain in three-dimensions. CLAMP de-compacts the genomic architecture of the X-chromosome by promoting long-range interactions and preventing the formation of shorter-range interactions (Fig. 4G). CLAMP not only tethers MSLc to the X-chromosome^20^ but also promotes long-range three-dimensional interactions involving MSLc binding sites (Fig. 4G). Previously, we demonstrated that CLAMP and MSLc function synergistically to increase transcript levels of X-linked genes^20,54^. Here, we show that both CLAMP and MSLc regulate the three-dimensional organization of the X-chromosome: CLAMP regulates three-dimensional interactions along the length of the entire X-chromosome while MSLc acts locally at several CES.

Throughout the genome, CLAMP promotes long-range interactions between active chromatin regions, including CES. However, we observe that CLAMP has a stronger function in mediating three-dimensional interactions on the X-chromosome than on autosomes. CLAMP is an ancient zinc-finger protein that is highly conserved across *Diptera* and is present on chromatin in the earliest stages of development before the more recently evolved MSLc^18,20,22,23^. Transposons containing CLAMP recognition sequences transposed onto the ancient X-chromosome and GA-repeats present at splice-junctions expanded, which together increased the density of CLAMP occupancy at CES^17,18^. The enhanced role of CLAMP on the X-chromosome compared with autosomes is likely due to the increased density of CLAMP binding sites within CES on the X-chromosome^18^.

Furthermore, CLAMP interacts with several insulator protein complexes^30,49^, which are known to modulate three-dimensional genomic interactions and contain CP190. We observe that the enrichment of insulator proteins is increased at sites where CLAMP functions to regulate threedimensional interactions, compared to non-interacting control regions. Therefore, it is likely that CLAMP regulates three-dimensional organization by modulating the ability of insulator proteins to drive the formation of three-dimensional interactions. For example, CLAMP is known to alter the occupancy of CP190, a component of several different insulator complexes^30^. It is also possible that insulator proteins alter the ability of CLAMP to regulate three-dimensional interactions.

Overall, we demonstrate that CLAMP, a transcription factor enriched on the male X-chromosome due to synergy with MSLc^20,21^, regulates long-range genomic interactions on the male X-chromosome including those involving CES. Therefore, CLAMP-mediated threedimensional interactions promote the formation of the three-dimensional active chromatin domain critical for dosage compensation. Further work will be required to define the functional relationships between CLAMP and all of the known insulator proteins.

## Acknowledgements

We thank Dr. Leila Rieder for critical reading of the manuscript and Dr. John Urban and Dr. Nicola Neretti for advice on computational analysis. This research was supported by NIH grant: 5R35GM126994. WJ was funded by a NSF GRFP Fellowship and HHMI Gilliam Fellowship.

## Author contributions

WJ performed all experiments, analysis and wrote the manuscript. EL provided advice on experiments and writing of the manuscript.

---

**Table S1. Processing of Hi-C and 4C sequencing reads.** This spreadsheet contains mapping and filtering data from processing Hi-C data with HiC-Pro and 4C-seq data with 4C-ker (see methods for analysis description).

**Table S2. 4C Viewpoints.** This spreadsheet contains the viewpoint coordinates, primers, and primary and secondary restriction enzymes.

**Table S3. High-confidence interactions.** This spreadsheet contains the high-confidence interactions identified for Hi-C and 4C-seq (see methods for analysis description).

**Table S4. Differential Analysis of Hi-C and 4C-seq.** This spreadsheet contains the results of differential analysis performed for Hi-C using diffHiC and 4C-seq using 4C-ker (see methods for analysis description).

## Methods

### Cell culture conditions

*Drosophila* S2 cells were maintained at 25C in Gibco Schneider’s *Drosophila* media (ThermoFisher Scientific) supplemented with 10% heat-inactivated fetal bovine serum (FBS) and 1.4X antibiotic-antimycotic (ThermoFisher Scientific). Cells were passaged every 2-3 days to maintain appropriate density.

### RNAi treatment

#### Generation of dsRNA for RNAi treatment

Generation of dsRNA targeting *gfp* (control), *clamp, msl2* and *trl* for RNAi have been previously validated and described^4,19,20,55^ PCR product was used as template to generate dsRNA with an ambion T7 MEGAscript kit (ThermoFisher Scientific); dsRNAs were purified following DNase treatment with a Qiagen RNeasy kit (Qiagen).

#### RNAi treatment

RNAi was performed in T75 tissue culture flasks. A total of 7×10^6 S2 cells were suspended in 6mL of Schneider’s *Drosophila* media (without FBS) and added to a T75 culture flask containing 67.5ug of *gfp, clamp, msl2,* or *trl* dsRNA suspended in 3mL of Invitrogen UltraPure water (ThermoFisher Scientific). Cells were serum starved for 45 minutes at room temperature, then 10.5mL of Schneider’s *Drosophila* media supplemented with 10% FBS was added. Cells were incubated for a total of 6 days as described previously^20,44^ After 6 days, samples were collected and RNAi knockdown was validated via western blotting or qRT-PCR. The remaining cells were collected by centrifugation and resuspended to 5×10^6 cells/ml in fresh non-FBS media. Fresh formaldehyde solution (36.5-38% in H_2_O; Sigma Aldrich) was then added to obtain a final concentration of approximately 1% formaldehyde. Cell suspensions were then incubated at RT on a rocking platform for 10 minutes; 2.5M glycine was then added (final concentration of 0.125M) to quench the formaldehyde; the cells were incubated as before for an additional 5 minutes. The cell suspension were then immediately placed on ice for 15 minutes. The cell suspension was then aliquoted into individual tubes (5 million cells per tube). The cell suspensions were centrifuged at 4C for 5 minutes at 3000 RPM, supernatant was removed and then flash frozen in liquid nitrogen and stored at −80C for future processing.

#### Knockdown validation

For Hi-C experiments knockdown of CLAMP was validated using the Western Breeze kit (Invitrogen). Antibodies used for detection were a previously described custom rabbit anti-CLAMP (1:1,000, Abcam)^22^. Mouse anti-actin (1:400,000, Sigma Aldrich) was used as a loading control.

#### Quantitative real-time PCR

To determine transcript abundance of *clamp, msl2, or trl* RNA was was calculated using the 2^-ΔΔCt^ method^56^ using RNA extracted from 500ul of cells following the 6 day incubation using an RNeasy Plus RNA extraction kit (Qiagen). Gene targets were amplified from cDNA using previously validated primers for *clamp, msl2, trl,* and three internal control genes *(gapdh, rpl32,* and *ras64b)* using triplicate technical replicates for each biological replicate per condition. Samples were normalized to the control *gfp* RNAi condition.

### Hi-C experimental procedure

Hi-C libraries of two independent biological replicates per RNAi condition *(clamp* and *gfp)* were generated as follows:

#### Cell lysis and restriction enzyme digestion

Approximately 10 million formaldehyde crosslinked (crosslinking procedure described above) S2 cells were resuspended in 500ul of fresh cold lysis buffer (10mM Tris-HCL pH 8.0, 10mM NaCl, 10ul of 50X Protease inhibitors cocktail, 0.2% Igepal CA630 in UltraPure water) and incubated on ice for 30 minutes. Cell suspensions were then centrifuged for 5 minutes at 3000 RPM at 4C and the supernatant removed. The cell pellet was resuspended with 300ul of cold 1x NEBuffer2. Cell suspensions were again centrifuged at 3000 RPM at 4C and the supernatant was removed. Cells were resuspended in 95ul of 1X NEBuffer2 and 5ul of 10% SDS was added. The cell suspension was homogenized by gentle pipetting and then incubated at 65C for 10 minutes. The cell suspension was then placed on ice and 200ul of 1X NEBuffer2 along with 60ul of 10% Triton X-100 (to quench the SDS) was added. The cell suspension was incubated at 37C for 15 minutes. Lysis efficiency and nuclei integrity was checked via microscope. The samples were centrifuged for 5 minutes at 3000 RPM at 4C and the supernatant was removed. The pellet was resuspended in 300ul of 1X NEBuffer2 and 400 U of HindIII restriction enzyme was added. The samples were incubated overnight at 37C. The following morning an additional 200U of HindIII was added to each of the samples and they were incubated an additional 2 hours at 37C.

#### End-repair, labeling, in-nuclei ligation, and crosslink reversal

The samples were centrifuged for at 3000 RPM for 5 minutes at 4C and the supernatant removed. The samples were resuspended with 250ul of 1X NEBuffer and 50ul of the following mix was added: 1.5ul of 10mM dATP, 1.5ul of 10mM dGTP, 1.5ul of 10mM dTTP, 37.5ul of 0.4mM Biotin-11-dCTP, 1ul of 50U/ul DNA Polymerase I Large (Klenow) fragment, and 7ul of UltraPure water. The sample were incubated at 37C for 45 minutes and then incubated at 65C for 15 minutes. The samples were centrifuged for 5 minutes at 3000 RPM and the supernatant discarded. Samples were resuspended in 1.195mL of the following ligation mix: 120ul of 10X T4 DNA Ligase Buffer, 100ul of 10% Triton X-100, 12ul of 10mg/mL BSA and 963ul of UltraPure water. 5ul of 2000U/ul T4 DNA ligase was then added to each sample and the samples were incubated at 16C overnight. The samples were centrifuged for 5 minutes at 3,000 RPM and supernatant removed. The samples were resuspended in 400ul of 1X NEBuffer2 and 10ul of 10mg/mL RNase A was added and the samples were incubated for 15 minutes at 37C at 300 RPM shaking. 20ul of 10mg/ml Proteinase K was added and the samples were incubated overnight at 65C to reverse crosslinks at 300 RPM shaking. The next morning an additional 20ul of Proteinase K was added again and the sample were incubated for 2 hours at 65C.

#### DNA purification

The samples were cooled to room temperature and 400ul of an equal volume of Phenol/Chloroform/Isoamyl Alcohol was added to each sample and the solution was mixed vigorously. The samples were centrifuged for 5 minutes at 13,000 RPM and the upper aqueous phase was transferred to a new tube. DNA was precipitated by adding 40ul of NaAc pH 5.2 and 1mL of 100% ethanol. The samples were incubated for 30 minutes at −80C and then centrifuged at 13,000 RPM for 30 minutes at 4C. The supernatant was discarded and pellet was washed with 1mL of Ethanol 70%. The sample was centrifuged for 15 minutes at 13,000 RPM at 4C. The supernatant was discarded and the pellet was air-dried for 5 minutes. The pellet was resuspended in 1X TE buffer.

DNA shearing and size selection, biotin pull-down and sequencing library preparation were performed as described in^36^. Multiplexed libraries were sequenced on an Illumina Hi-Seq 2500 configured for 150-bp paired-end reads.

### 4C-seq experimental procedure

4C-seq libraries were generated from S2 cells. Nuclear extraction and crosslinking were carried out as described in Hi-C experimental procedure above. The remainder of the 4C-seq procedure was carried out as previously described^57^. Csp6I, DpnII or NlaIII were used as primary or secondary restriction enzymes. Primer sequences for each viewpoint and condition were generated using 4C primer design (https://mnlab.uchicago.edu/4Cpd/) and are listed in supplemental table S2. Independent biological replicates of multiplexed libraries were sequenced on separate lanes of an Illumina Hi-Seq 2500 configured for 150-bp paired-end reads.

### Datasets

CES locations were obtained from GSE39271. Hi-C following RNAi against *msl2, msl3,* or *gfp* was obtained from GSE58821. MACC data was obtained from GSE99894. S2 cell chip-seq data sets for CP190, SuHw, Mod, and dCTCF GSE41354. S2 cell chip-seq data sets for GAF and CLAMP were obtained from GSE107059.

Raw and processed sequencing data generated is deposited to NCBI GEO under accession GSE130546.

### Hi-C analysis

Hi-C data was processed using the dm6 *Drosophila* reference genome^58^ using HiC-Pro pipeline version 2.7.8^59^ for read mapping (MIN_MAPQ = 15), filtering and quality checks to generate valid read pairs and interaction matrices. GenomeDisco version 1.0.0^38^ was utilized to determine concordance between biological replicates. Custom scripts were used to convert Hi-C-Pro interaction matrices to a format suitable for input into GenomeDisco. For matrix visualizations valid pairs were processed into .hic files using the hicpro2juicebox script provided with HiC-Pro. Matrices were visualized using Juicebox^60^ with KR matrix balancing.

#### PCA and DLR analysis

Valid pairs for biological replicates were merged and imported into Homer version 4.10^61^ using the command makeTagDirectory with the parameter -format HiCsummary. Principal component analysis (PCA) was performed using Homer with the command runHiCpca.pl at resolutions of 10kb, 20kb and 50kb. Distal to local log2 ratio analysis was performed using Homer with the command analyzeHiC with the following parameters: -res 5000 -window 15000 – compactionStats auto -dlrDistance 250000. Distal to local log2 ratio differential analysis was performed using the command subtractBedGraphsDirectory.pl with the parameter -center.

#### Identification of high confidence long-range interactions

20kb resolution contact matrices corresponding to the control *(gfp* RNAi) conditions were generated and iterative corrected using HiC-Pro. The resultant contact matrices were then adapted for Fit-Hi-C version 2.0.5^40^ using the hicpro2Fit-Hi-C.py script provided with Hi-C-Pro. Raw contact matrices and ICE biases were used as input into Fit-Hi-C to identify significant contacts with the following parameters: -L 5000000 -U 15000000 -v -b 100 -p2. Significant contacts were filtered for those with q < 0.01 and the intersection of significant contacts (q < 0.01) from the replicates was used as the high confidence interaction set.

#### Identifying Differential interactions (DIs)

diffHiC version 1.14.0^41^ which uses the edgeR^42^ package for differential statistics was utilized to identify differential interactions. Valid pairs for from HiC-Pro for each replicate was imported into diffHic using the savePairs function. Low-abundance read pairs were filtered out and the resulting data was normalized with TMM normalization and trended biases were removed. Differential interactions were identified using a bin size of 30kb, an FDR target of 0.05, and the functions diClusters, combineTests, and getBestTest. No threshold was applied for log ratio.

### 4C-Seq analysis

4C sequencing reads were demultiplexed based on index sequences and inline barcodes. Sequencing reads corresponding to the reading primer were aligned to a reduced genome of unique sequences adjacent to restriction enzyme sites derived from the dm6 *Drosophila* reference genome^58^ using Bowtie2 version 2.3.0^62^ with the following parameters: -N 0 −5 24 −3 101 --very-sensitive, which removes barcode and primer sequences and trim the read length to 25bp prior to mapping. The aligned reads were then processed as described for use with the 4C-Ker pipeline^51^ which uses a three state Hidden-Markov Model to find chromosome-wide interactions and DESeq2^63^ to perform differential analysis. Quality checks were performed within the 4C-Ker pipeline. *Cis* analysis was performed for each viewpoint using k=10 and p-value cutoff of 0.05 was used for differential analysis. Viewpoints were visualized using IGV^64^

#### Identification of high confidence long-range interactions

A set of high confidence interactions for each viewpoint was determined by intersecting the significant interactions individually detected in each replicate of the control *(gfp* RNAi) condition using BEDTools version 2.27.1^45^.

### Comparison of 4C and Hi-C high confidence interactions

High confidence Hi-C interactions involving each 4C-seq viewpoint was determined by intersecting the viewpoint coordinate sequence with the Hi-C high confidence interactions set. This set of interactions was then intersected with the 4C-seq high confidence interactions set to determine overlapping pairwise high confidence interactions identified in both experiments. All intersections were performed using BEDTools.

### Chromatin state annotation

Chromatin states were from the modENCODE project (DCC id: modENCODE_33lifted over to dm6, using USC6S3)^43,65^ and were lifted over to dm6 with the USCS liftOver tool (http://genome.ucsc.edu)^66^. Ratios were normalized to account for the per chromosome abundance of each individual chromatin state. Jaccard similarity scores were computed using BEDTools.

### Distance calculations

Distances to nearest CES were calculated using BEDTools. Controls were conducting by 100 permutations of randomly shuffling CES restricting reshuffled regions to either active or inactive regions as defined by chromatin state^43^.

### Chromatin accessibility (MACC) analysis

MACC data generated following RNAi knockdown of *gfp* (control), *clamp,* or *msl2* was lifted over to dm6 with the USCS liftover tool. MACC data was intersected with genomic regions of interest using BEDTools to determine corresponding MACC scores. Other analyses were performed in the python programming environment (https://www.python.org).

### Chip-seq data processing and generation of high confidence peak sets

For each ChIP-seq dataset used, the respective sequencing reads were downloaded and mapped to release 6 *D. melanogaster* genome (dm6)^58^ using Bowtie2 with parameter -N 1. Reads with a MAPQ <30 and PCR duplicate reads identified using Picard MarkDuplicates version 2.9.2^67^ were removed using SAMtools version 1.9^68^. Reads mapped to dm3 blacklisted regions (https://sites.google.com/site/anshulkundaje/projects/blacklists) lifted over to dm6, using USCS liftOver, were also removed. In cases where biological replicates were not available the aligned reads were split into pseudoreplicates. MACS2 version 2.1.1^69^ was used to identify peaks with the following parameters: --nomodel -B --SPMR --keep-dup all -f AUTO -g dm -p 0.01. MACS2 was also used generate fold enrichment (default parameters) bedGraphs for each factor. In order to reduce the number of false positive peaks the irreproducible discovery rate (IDR) was calculated using IDR version 2.0.3^70^ using the MACS2 peak score calculated for each replicate experiment (biological replicate in the case of GAF and CLAMP; pseudoreplicate in the case of CP190, SuHw, Mod, dCTCF) as input to IDR. Peaks with an IDR < 0.01 were retained (In cases where there were more than two biological replicates pairwise IDR comparison of each replicate was made and the longest resulting peak list was used). BEDTools and USCS bedGraphToBigWig^71^ tool was used to convert bedGraphs to bigwig format. Average profiles were generated using deepTools version 3.1.0^72^.

## Supplemental Figure Legends

**Figure S1.**
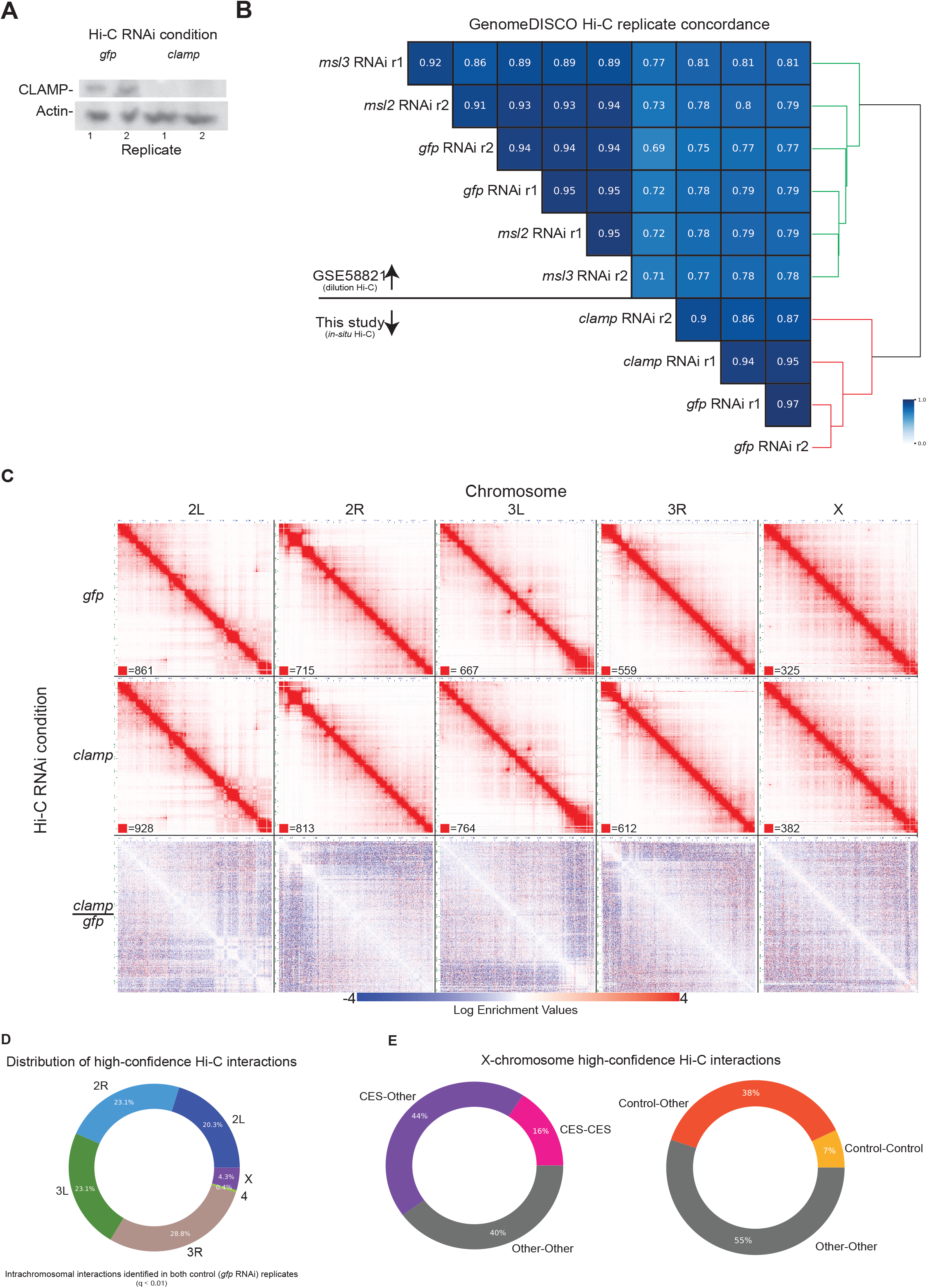
Experimental QC and Hi-C Replicate Concordance. A. Western blot for *gfp* and *clamp* RNAi Hi-C experiments indicating successful knockdown of CLAMP protein (Source data provided as a Source Data file). B. Pairwise Hi-C replicate concordance scores as measured by GenomeDISCO (Source data provided as a Source Data file). C. Per chromosome KR balanced matrices for *gfp* RNAi (top), *clamp* RNAi (middle), and *clamp* vs *gfp* RNAi (bottom). Each chromosome is shown at 50 kb resolution. D. The per chromosome distribution of high confidence Hi-C interactions (Source data provided in Table S3 and Source Data file). E. The distribution of X-chromosome high confidence Hi-C interactions that involve a CES and matched control (randomized active chromatin regions) (Source data provided as a Source Data file).

**Figure S2.**
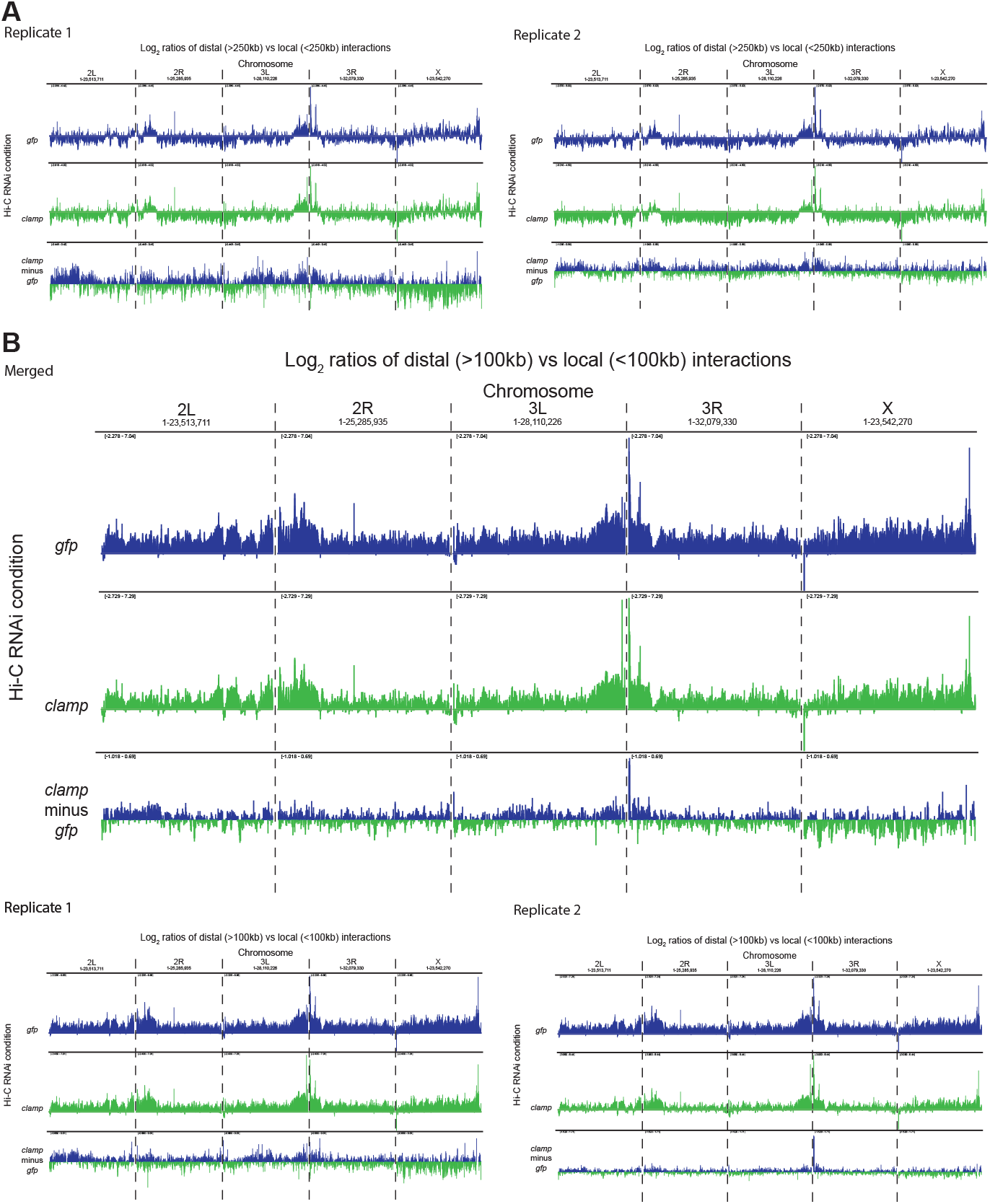
CLAMP regulates the length span of genomic interactions on the male X-chromosome, related to Figure 1. **A.** Per chromosome distal vs local ratio (DLR) for *gfp* RNAi (blue), *clamp* RNAi (green), and *clamp* vs *gfp* RNAi (bottom). For the *clamp* vs *gfp* RNAi comparison, a positive number (blue) indicates the ratio of distal vs. local interactions becomes higher following *clamp* RNAi. A negative number (green) indicates the ratio of distal vs. local interactions becomes lower following *clamp* RNAi. Similar to Figure 1. but shown for paired individual replicates. **B.** Per chromosome (DLR) for merged (top) and paired individual replicates (bottom) using a distance > 100kb to denote distal interactions.

**Figure S3.**
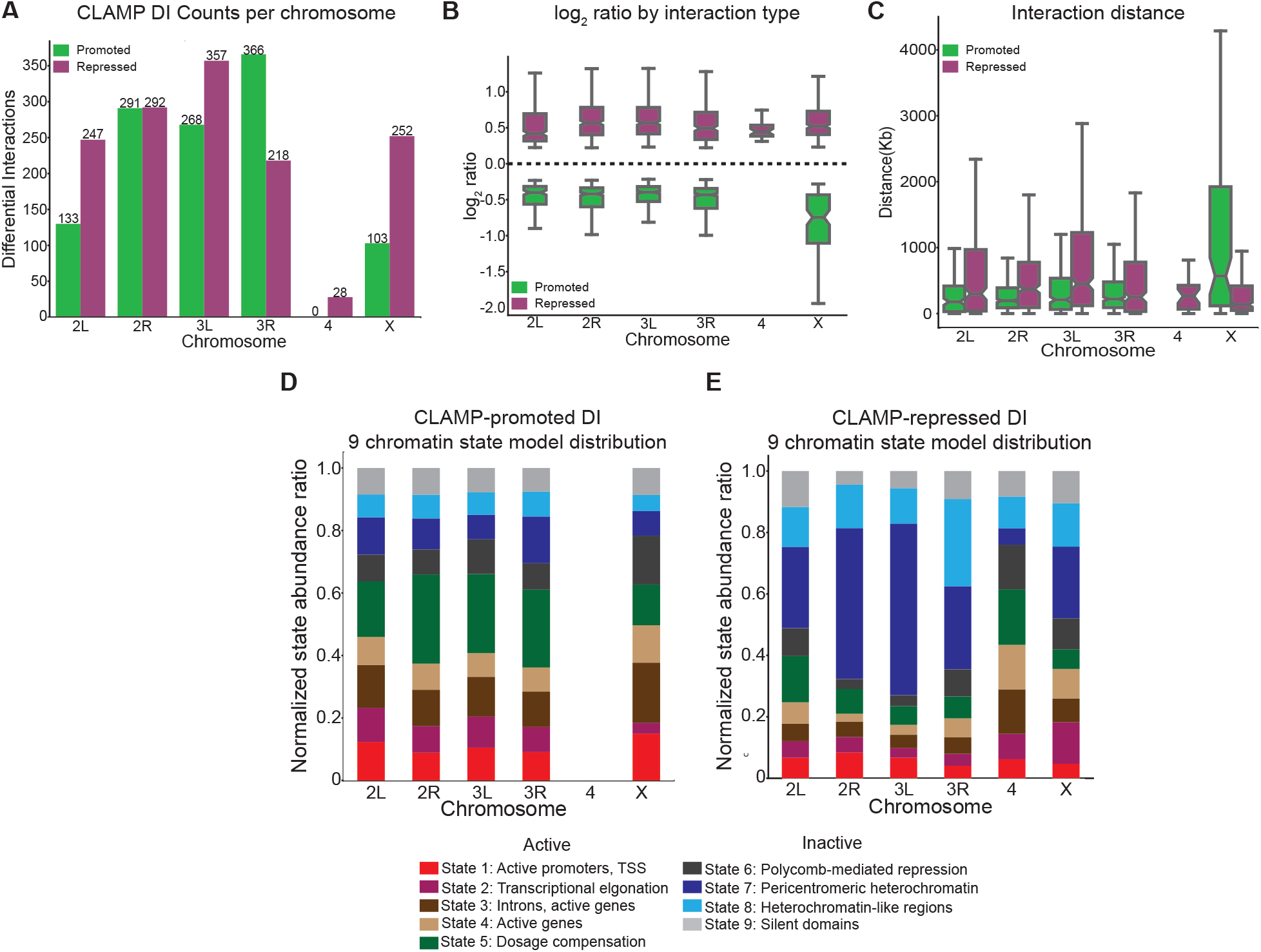
CLAMP promotes long-range interactions on the X-chromosome and generally promotes interactions in active chromatin and represses interactions in inactive chromatin. A. Per chromosome differential interaction count of CLAMP-promoted and CLAMP-repressed interactions (Source data provided in Table S4). B. Per chromosome log2 ratios by interaction type for CLAMP-promoted and CLAMP-repressed interactions, related to Figure 1A (Source data provided in Table S4). C. Per chromosome distribution of distances between differential interaction anchors for CLAMP-promoted and CLAMP-repressed interactions, related to Figure 1B (Source data provided in Table S4). D. Per chromosome normalized ratio of chromatin states occurring at CLAMP-promoted interactions (Source data provided as a Source Data file). E. Per chromosome normalized ratio of chromatin states occurring at CLAMP-repressed interactions (Source data provided as a Source Data file). For all box and whisker plots, the 95% confidence interval is shown with a notch around the median line; whiskers represent 1.5 IQR; outliers have been omitted.

**Figure S4.**
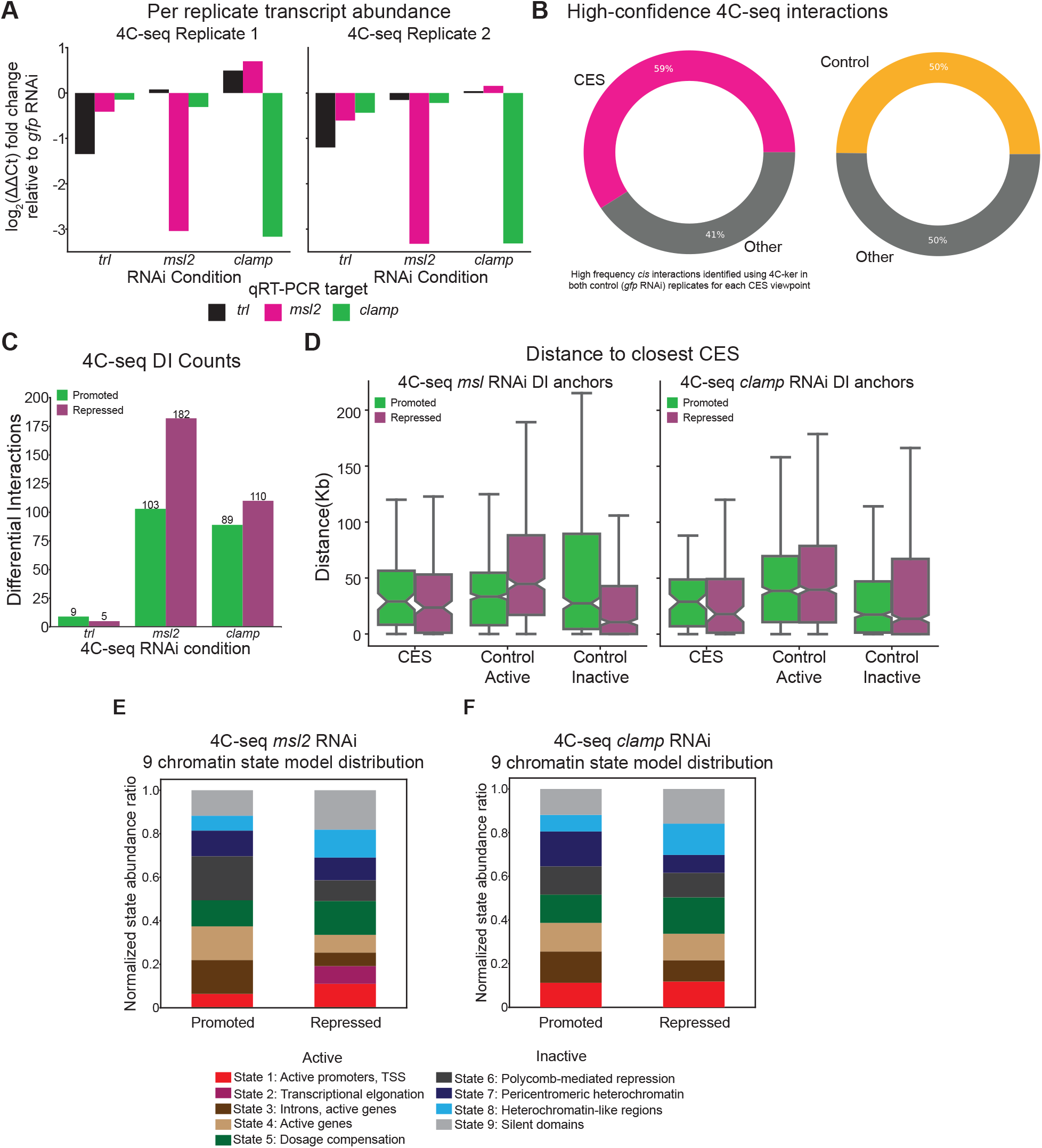
Summary of high-resolution 4C-seq analysis. A. Quantitative real-time PCR indicates successful RNAi knockdown of each target gene. Plotted is the log_2_ fold change (ΔΔCt) for each biological replicate after internal normalization to three control genes *(gapdh, rpl32,* and *ras64b).* Samples are normalized to the *gfp* RNAi condition (Source data provided as a Source Data file). B. Percentage of high-confidence *cis* interactions for all 4C-seq viewpoints (control *gfp* RNAi condition) that correspond to a region containing a CES and matched control (randomized active chromatin regions) (Source data provided as a Source Data file). C. 4C-seq differential interaction counts per RNAi condition identified using 4C-ker (Source data provided in Table S4). D. Distribution of distances to nearest CES or control CES for MSL2 and CLAMP-promoted and CLAMP-repressed interactions. For all box and whisker plots, the 95% confidence interval is shown with a notch around the median line; whiskers represent 1.5 IQR, outliers have been omitted (for *msl* RNAi anchors: Promoted n = 103, repressed n = 182, for *clamp* RNAi anchors: Promoted n = 89, repressed n = 110; distribution of controls obtained by 100 permutations of randomly shuffling CES (see methods; Source data provided as a Source Data file). E. Ratio of chromatin states, normalized for X-chromosome state abundance, occurring at MSL2-promoted and MSL2-repressed interactions (Source data provided as a Source Data file). F. Ratio of chromatin states, normalized for X-chromosome state abundance, occurring at CLAMP-promoted and CLAMP-repressed interactions (Source data provided as a Source Data file).

## References

1. Strom, A. R. et al. Phase separation drives heterochromatin domain formation. Nature 547, 241–245 (2017).

2. Sexton, T. & Cavalli, G. The role of chromosome domains in shaping the functional genome. Cell 160, 1049–1059 (2015).

3. Ghavi-Helm, Y. et al. Highly rearranged chromosomes reveal uncoupling between genome topology and gene expression. Nat. Genet. 51, 1272–1282 (2019).

4. Hamada, F. N., Park, P. J., Gordadze, P. R. & Kuroda, M. I. Global regulation of X chromosomal genes by the MSL complex in Drosophila melanogaster. Genes Dev. 19, 2289–2294 (2005).

5. Conrad, T. et al. The MOF Chromobarrel Domain Controls Genome-wide H4K16 Acetylation and Spreading of the MSL Complex. Dev. Cell 22, 610–624 (2012).

6. Straub, T. & Becker, P. B. Dosage compensation: The beginning and end of generalization. Nat. Rev. Genet. 8, 47–57 (2007).

7. Jordan, W., Rieder, L. E. & Larschan, E. Diverse Genome Topologies Characterize Dosage Compensation across Species. Trends Genet. 35, 308–315 (2019).

8. Belote, J. M. & Lucchesi, J. C. Male-specific lethal mutations of Drosophila melanogaster. Genetics 96, 165–186 (1980).

9. Lucchesi, J. C. Dosage compensation in Drosophila and the ‘complex’ world of transcriptional regulation. BioEssays 18, 541–547 (1996).

10. Kuroda, M. I., Kernan, M. J., Kreber, R., Ganetzky, B. & Baker, B. S. The maleless protein associates with the X chromosome to regulate dosage compensation in drosophila. Cell 66, 935–947 (1991).

11. Hilfiker, A., Hilfiker-Kleiner, D., Pannuti, A. & Lucchesi, J. C. mof, a putative acetyl transferase gene related to the Tip60 and MOZ human genes and to the SAS genes of yeast, is required for dosage compensation in Drosophila. EMBO J. 16, 2054–2060 (1997).

12. Meller, V. H. & Rattner, B. P. The roX genes encode redundant male-specific lethal transcripts required for targeting of the MSL complex. EMBO J. 21, 1084–1091 (2002).

13. Franke, A. & Baker, B. S. The rox1 and rox2 RNAs are essential components of the compensasome, which mediates dosage compensation in Drosophila. Mol. Cell 4, 117–122 (1999).

14. Alekseyenko, A. A. et al. A Sequence Motif within Chromatin Entry Sites Directs MSL Establishment on the Drosophila X Chromosome. Cell 134, 599–609 (2008).

15. Kelley, R. L. et al. Epigenetic spreading of the Drosophila dosage compensation complex from roX RNA genes into flanking chromatin. Cell 98, 513–522 (1999).

16. Straub, T., Grimaud, C., Gilfillan, G. D., Mitterweger, A. & Becker, P. B. The chromosomal high-affinity binding sites for the Drosophila dosage compensation complex. PLoS Genet. 4, 1–14 (2008).

17. Ellison, C. & Bachtrog, D. Contingency in the convergent evolution of a regulatory network: Dosage compensation in drosophila. PLoS Biol. 17, e3000094 (2019).

18. Kuzu, G. et al. Expansion of GA dinucleotide repeats increases the density of CLAMP binding sites on the X-chromosome to promote Drosophila dosage compensation. PLoS Genet. 12, e1006120 (2016).

19. Larschan, E. et al. Identification of chromatin-associated regulators of MSL complex targeting in Drosophila dosage compensation. PLoS Genet. 8, (2012).

20. Soruco, M. M. L. et al. The CLAMP protein links the MSL complex to the X chromosome during Drosophila dosage compensation. Genes Dev. 27, 1551–1556 (2013).

21. Albig, C. et al. Factor cooperation for chromosome discrimination in Drosophila. Nucleic Acids Res. 47, 1706–1724 (2019).

22. Rieder, L. E. et al. Histone locus regulation by the Drosophila dosage compensation adaptor protein CLAMP. Genes Dev. 31, 1494–1508 (2017).

23. Franke, A., Dernburg, A., Bashaw, G. J. & Baker, B. S. Evidence that MSL-mediated dosage compensation in Drosophila begins at blastoderm. Development 122, 2751–2760 (1996).

24. Lott, S. E. et al. Noncanonical compensation of zygotic X transcription in early Drosophila melanogaster development revealed through single-embryo RNA-Seq. PLoS Biol. 9, e1000590 (2011).

25. Urban, J. et al. Enhanced chromatin accessibility of the dosage compensated Drosophila male X-chromosome requires the CLAMP zinc finger protein. PLoS One 12, e0186855 (2017).

26. Alekseyenko, A. A., Larschan, E., Lai, W. R., Park, P. J. & Kuroda, M. I. High-resolution ChIP-chip analysis reveals that the Drosophila MSL complex selectively identifies active genes on the male X chromosome. Genes Dev. 20, 848–857 (2006).

27. Larschan, E. et al. X chromosome dosage compensation via enhanced transcriptional elongation in Drosophila. Nature 471, 115–118 (2011).

28. Ramírez, F. et al. High-Affinity Sites Form an Interaction Network to Facilitate Spreading of the MSL Complex across the X Chromosome in Drosophila. Mol. Cell 60, 146–162 (2015).

29. Schauer, T. et al. Chromosome topology guides the Drosophila Dosage Compensation Complex for target gene activation. EMBO Rep. 18, 1854–1868 (2017).

30. Bag, I., Dale, R. K., Palmer, C. & Lei, E. P. The zinc-finger protein CLAMP promotes gypsy chromatin insulator function in Drosophila. J. Cell Sci. 132, jcs-226092 (2019).

31. Darrow, E. M. et al. Deletion of DXZ4 on the human inactive X chromosome alters higher-order genome architecture. Proc. Natl. Acad. Sci. U. S. A. 113, E4504–E4512 (2016).

32. Kieffer-Kwon, K. R. et al. Myc Regulates Chromatin Decompaction and Nuclear Architecture during B Cell Activation. Mol. Cell 67, 566–578.e10 (2017).

33. Muller, M., Hagstrom, K., Gyurkovics, H., Pirrotta, V. & Schedl, P. The Mcp element from the Drosophila melanogaster bithorax complex mediates long-distance regulatory interactions. Genetics 153, 1333–1356 (1999).

34. Noordermeer, D. et al. The dynamic architecture of Hox gene clusters. Science (80-.). 334, 222–225 (2011).

35. Vian, L. et al. The Energetics and Physiological Impact of Cohesin Extrusion. Cell 173, 1165–1178.e20 (2018).

36. Rao, S. S. P. et al. A 3D map of the human genome at kilobase resolution reveals principles of chromatin looping. Cell 159, 1665–1680 (2014).

37. Schneider, I. Cell lines derived from late embryonic stages of Drosophila melanogaster. J. Embryol. Exp. Morphol. 27, 353–365 (1972).

38. Ursu, O. et al. GenomeDISCO: A concordance score for chromosome conformation capture experiments using random walks on contact map graphs. Bioinformatics 34, 2701–2707 (2018).

39. Heinz, S. et al. Transcription Elongation Can Affect Genome 3D Structure. Cell 174, 1522–1536.e22 (2018).

40. Ay, F., Bailey, T. L. & Noble, W. S. Statistical confidence estimation for Hi-C data reveals regulatory chromatin contacts. Genome Res. 24, 999–1011 (2014).

41. Lun, A. T. L. & Smyth, G. K. diffHic: A Bioconductor package to detect differential genomic interactions in Hi-C data. BMC Bioinformatics 16, 258 (2015).

42. Robinson, M. D., McCarthy, D. J. & Smyth, G. K. edgeR: A Bioconductor package for differential expression analysis of digital gene expression data. Bioinformatics 26, 139–140 (2009).

43. Kharchenko, P. V et al. Comprehensive analysis of the chromatin landscape in Drosophila melanogaster. Nature 471, 480–5 (2011).

44. Kaye, E. G. et al. Differential Occupancy of Two GA-Binding Proteins Promotes Targeting of the Drosophila Dosage Compensation Complex to the Male X Chromosome. Cell Rep. 22, 3227–3239 (2018).

45. Quinlan, A. R. BEDTools: The Swiss-Army tool for genome feature analysis. Curr. Protoc. Bioinforma. 2014, 11.12.1–11.12.34 (2014).

46. Grimaud, C. & Becker, P. B. The dosage compensation complex shapes the conformation of the X chromosome in Drosophila. Genes Dev. 23, 2490–2495 (2009).

47. Farkas, G. et al. The Trithorax-like gene encodes the Drosophila GAGA factor. Nature 371, 806–808 (1994).

48. Kasinathan, S., Orsi, G. A., Zentner, G. E., Ahmad, K. & Henikoff, S. High-resolution mapping of transcription factor binding sites on native chromatin. Nat. Methods 11, 203–209 (2014).

49. Kaye, E. G. et al. Drosophila Dosage Compensation Loci Associate with a Boundary-Forming Insulator Complex. Mol. Cell. Biol. 37, MCB–00253 (2017).

50. Greenberg, A. J., Yanowitz, J. L. & Schedl, P. The Drosophila GAGA Factor is Required for Dosage Compensation in Males and for the Formation of the Male-Specific-Lethal Complex Chromatin Entry Site at 12DE. Genetics 166, 279–289 (2004).

51. Raviram, R. et al. 4C-ker: A Method to Reproducibly Identify Genome-Wide Interactions Captured by 4C-Seq Experiments. PLoS Comput. Biol. 12, e1004780 (2016).

52. Gavrilov, A. A. et al. Disclosure of a structural milieu for the proximity ligation reveals the elusive nature of an active chromatin hub. Nucleic Acids Res. 41, 3563–3575 (2013).

53. Van De Werken, H. J. G. et al. Robust 4C-seq data analysis to screen for regulatory DNA interactions. Nat. Methods 9, 969–972 (2012).

54. Urban, J. A., Urban, J. M., Kuzu, G. & Larschan, E. N. The Drosophila CLAMP protein associates with diverse proteins on chromatin. PLoS One 12, e0189772 (2017).

55. Fuda, N. J. et al. GAGA Factor Maintains Nucleosome-Free Regions and Has a Role in RNA Polymerase II Recruitment to Promoters. PLOS Genet. 11, e1005108 (2015).

56. Livak, K. J. & Schmittgen, T. D. Analysis of relative gene expression data using real-time quantitative PCR and the 2-A\CT method. Methods 25, 402–408 (2001).

57. Ghavi-Helm, Y. et al. Enhancer loops appear stable during development and are associated with paused polymerase. Nature 512, 96–100 (2014).

58. Hoskins, R. A. et al. The Release 6 reference sequence of the Drosophila melanogaster genome. Genome Res. 25, 445–458 (2015).

59. Servant, N. et al. HiC-Pro: An optimized and flexible pipeline for Hi-C data processing. Genome Biol. 16, 259 (2015).

60. Durand, N. C. et al. Juicebox Provides a Visualization System for Hi-C Contact Maps with Unlimited Zoom. Cell Syst. 3, 99–101 (2016).

61. Heinz, S. et al. Simple Combinations of Lineage-Determining Transcription Factors Prime cis-Regulatory Elements Required for Macrophage and B Cell Identities. Mol. Cell 38, 576–589 (2010).

62. Langmead, B. & Salzberg, S. L. Fast gapped-read alignment with Bowtie 2. Nat. Methods 9, 357–359 (2012).

63. Love, M. I., Huber, W. & Anders, S. Moderated estimation of fold change and dispersion for RNA-seq data with DESeq2. Genome Biol. 15, 550 (2014).

64. Thorvaldsdóttir, H., Robinson, J. T. & Mesirov, J. P. Integrative Genomics Viewer (IGV): High-performance genomics data visualization and exploration. Brief. Bioinform. 14, 178–192 (2013).

65. Roy, S. et al. Identification of Functional Elements and Regulatory Circuits by Drosophila modENCODE. Science (80-.). 330, 1787–1797 (2016).

66. James Kent, W. et al. The human genome browser at UCSC. Genome Res. 12, 996–1006 (2002).

67. Van der Auwera, G. A. et al. From fastQ data to high-confidence variant calls: The genome analysis toolkit best practices pipeline. in Current Protocols in Bioinformatics 43, 11.10.1–11.10.33 (John Wiley & Sons, Inc., 2013).

68. Li, H. et al. The Sequence Alignment/Map format and SAMtools. Bioinformatics 25, 2078–2079 (2009).

69. Zhang, Y. et al. Model-based analysis of ChIP-Seq (MACS). Genome Biol. 9, R137 (2008).

70. Li, Q., Brown, J. B., Huang, H. & Bickel, P. J. Measuring reproducibility of high-throughput experiments. Ann. Appl. Stat. 5, 1752–1779 (2011).

71. Kent, W. J., Zweig, A. S., Barber, G., Hinrichs, A. S. & Karolchik, D. BigWig and BigBed: Enabling browsing of large distributed datasets. Bioinformatics 26, 2204–2207 (2010).

72. Ramírez, F., Dündar, F., Diehl, S., Grüning, B. A. & Manke, T. DeepTools: A flexible platform for exploring deep-sequencing data. Nucleic Acids Res. 42, W187–W191 (2014).

